# Cryo-EM structure of the circumsporozoite protein of *Plasmodium falciparum* with a vaccine-elicited antibody reveals maturation of inter-antibody contacts

**DOI:** 10.1101/332304

**Authors:** David Oyen, Jonathan L. Torres, Christopher A. Cottrell, C. Richter King, Ian A. Wilson, Andrew B. Ward

**Affiliations:** Department of Integrative Structural and Computational Biology, The Scripps Research Institute, La Jolla, CA, USA; PATH’s Malaria Vaccine Initiative, PATH’s Center for Vaccine Innovation and Access, Washington, DC 20001, USA; The Skaggs Institute for Chemical Biology, The Scripps Research Institute, La Jolla, CA, USA

## Abstract

The circumsporozoite protein (CSP) on the surface of *Plasmodium falciparum* sporozoites is important for parasite development, motility, and host hepatocyte invasion. However, intrinsic disorder of the NANP repeat sequence in the central region of CSP has hindered its structural and functional characterization. Here, the cryo-EM structure at ∼3.4 Å resolution of a recombinant shortened CSP construct (rsCSP) with the variable domains (Fabs) of a highly protective monoclonal antibody reveals an extended spiral conformation of the central NANP repeat region surrounded by antibodies. This unusual structure appears to be stabilized and/or induced by interaction with an antibody where contacts between adjacent Fabs are somatically mutated and enhance the interaction. Such maturation in non-antigen contact residues may be an effective mechanism for antibodies to target tandem repeat sequences and provide novel insights into malaria vaccine design.

**Summary:** An unusual spiral conformation is formed for the NANP repeat region in *Plasmodium falciparum* circumsporozoite protein (CSP) in complex with antibodies generated by the RTS,S vaccine and is stabilized by affinity-matured inter-Fab interactions.

---

With an estimated 445,000 deaths and 216 million cases in 2016, malaria poses a significant threat to public health (1). Emerging resistance against current front-line anti-malarials and insecticide resistance has furthered the need for an efficient malaria vaccine candidate (2). The pre-erythrocytic stage of the *Plasmodium falciparum* life cycle is an ideal target for the development of a vaccine that breaks the cycle of infection. After a bite from an infected mosquito, *P. falciparum* sporozoites (PfSPZs) migrate from the skin to the hepatocytes. Immunization with irradiated PfSPZs can induce strong protective immune responses in mice, monkeys and humans (3). For many years, the leading target for vaccine design has been the major surface protein of sporozoites, the circumsporozoite protein (PfCSP), which contains a central region consisting of multiple NANP repeats (4) that vary (from 25 to 49) among different *P. falciparum* isolates (5, 6). In addition, PfCSP contains a flexible N-terminal domain with a heparin sulfate binding site for hepatocyte attachment (7) and a structured C-terminal domain with a thrombospondin-like type I repeat (*α,*TSR) (8). The most advanced malaria vaccine to date is RTS,S, formulated in GlaxoSmithKline‘s adjuvant AS01. RST,S contains part of PfCSP, including 19 NANP repeats and the *ケ*TSR domain, fused with hepatitis B surface antigen (HBsAg) (Fig. 1E) such that virus-like particles are formed when co-expressed with free HBsAg in yeast (9). The RTS,S vaccine has been shown to confer reasonable protection in children against clinical malaria (5-17 months old) with 51% protection over the first year of follow-up after a 0,1,2 month vaccination schedule [95% CI 48-55%]. Efficacy was see to wane to 26% over a 48 month follow up period [95% CI 21-31%]; however if a fourth boost was administered at month 20 post vaccination, efficacy is 39% [95% CI 34-43%]. (10-12). These results indicate that while the RTS,S vaccine is promising, an important objective in current malaria research is to improve and extend the vaccine efficacy. The R21 vaccine is an alternative approach to PfCSP with is composed of a single subunit with the same region of PfCSP fused to HBsAg (13). Clinical testing is underway using Matrix-M adjuvant as well as AS01.

**Fig. 1.**
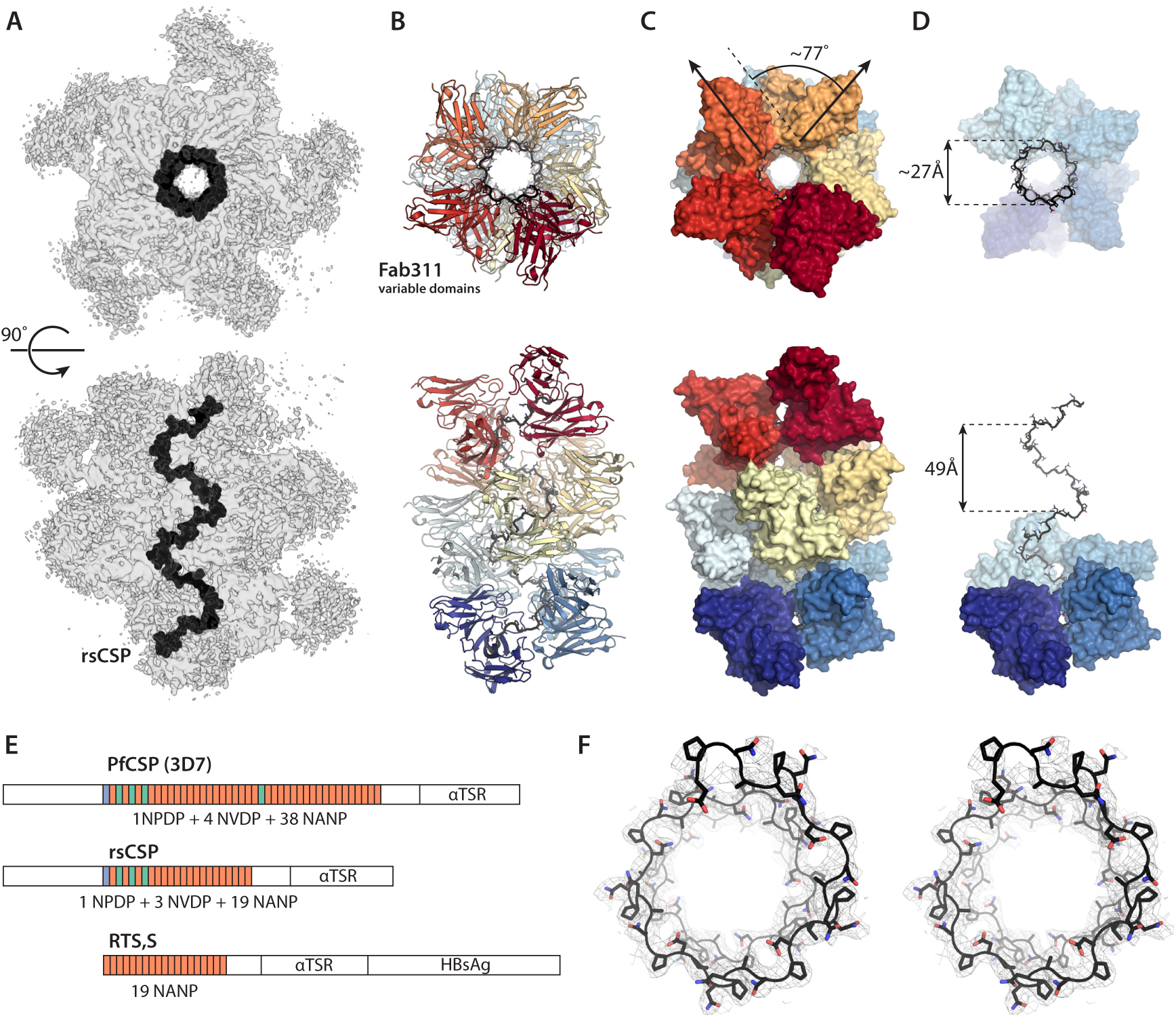
Spiral architecture of the rsCSP-Fab311 complex. (**A**) Side and top views of the cryo-EM electron density map are shown as gray transparent surfaces. The density for the Fab constant domains is weak due to flexibility around the elbow angle. rsCSP is colored in black. (**B**) The variable domains for Fab311 are built in the cryo-EM map and shown in ribbon representation. The Fabs are colored ranging from dark red to dark blue. (**C**) The rsCSP-Fab311 structure is shown as a molecular surface to visualize the orientation of the Fabs. Each individual Fab variable domain is rotated ∼77° with respect to the corresponding variable domain in a neighboring Fab. (**D**) Seven consecutive Fabs are stripped from the structure to unveil the buried repeat region of rsCSP which adopts a spiral. The pitch and diameter of the rsCSP spiral are 49 Å and ∼27 Å, respectively. (**E**) Schematic diagrams of PfCSP, rsCSP and RTS,S show the organization of the repeats within these three protein constructs. (F) Stereo representation of the rsCSP spiral (black carbons) and the cryo-EM density contoured at 5σ as a gray mesh.

One approach to improve vaccine designs, increasing the potency of antibody responses, involves investigation of the monoclonal antibody (mAb) responses generated through either whole PfSPZ or RTS,S immunization at the structural and functional level Recent X-ray structures of protective human fragments antigen-binding (Fabs) in complex with PfCSP repeat peptides revealed similarities and differences in how these repeats are recognized (14-17). Namely, the peptides are organized into NPNA structural units that can adopt type I β-turns and pseudo 3_10_ turns as originally observed for free peptides in solution and in peptide crystal structures (18, 19). One of these antibodies, mAb311, was isolated from a protected volunteer in a phase IIa RTS,S/AS01B controlled human malaria infection (CHMI) clinical trial (20) and inhibited parasite development in the liver by ∼97% as assessed by mouse challenge experiments with engineered *P. berghei* SPZs that express PfCSP (14). Interestingly, a low-resolution negative-stain electron microscopy (nsEM) reconstruction of a recombinant shortened PfCSP construct (rsCSP, Fig. 1E) in complex with Fabs of mAb311 (Fab311), gave the first insight into organization of the NANP repeats with bound antibodies. However, a high-resolution structure would provide valuable information for optimal display of protective epitopes in a vaccine setting.

## Cryo-EM structure of CSP and architecture of rsCSP-Fab311 complex

To decipher the architecture of the rsCSP-Fab311 complex at high resolution, we used single particle cryo-electron microscopy (cryo-EM). A final dataset of 206,991 particles was refined asymmetrically, resulting in an ∼3.4 Å resolution reconstruction (fig. S1). Eleven copies of the crystal structure of Fab311-(NPNA) _3_ could be fit into the EM map and the rsCSP-peptide complex was then assembled in COOT to generate an initial model. This model was subjected to multiple rounds of refinement into the EM density map using RosettaRelax (Fig. 1A, fig. S1 and S2, table S1).

The repeat region of rsCSP is well defined with continuous cryo-EM density (Fig. 1F and Fig. 2C) and forms an unusual extended spiral structure (Fig. 1A and D), from which multiple Fab311 antibodies radiate tangentially in a pseudo-helical arrangement (Fig. 1B), consistent with a previous negative-stain EM reconstruction (14). In the cryo-EM map, however, two additional Fabs were observed, demonstrating that 11 Fabs can bind simultaneously to rsCSP (Fig. 1C), although the density for the N- and C-terminal Fabs were sparse (table S1). In addition, no density was observed for the N-terminal or C-terminal αTSR domains of rsCSP, likely due to flexibility. Even though the αTSR domain has been observed to be structured by itself (8), it is connected to the NANP repeats through a disordered linker that is devoid of epitopes for Fab311. The angular twist between Fab variable domains is ∼77° with respect to each other, where 4.7 Fabs (360°/77°) are required to complete one full turn of the spiral (Fig. 1C).

**Fig. 2.**
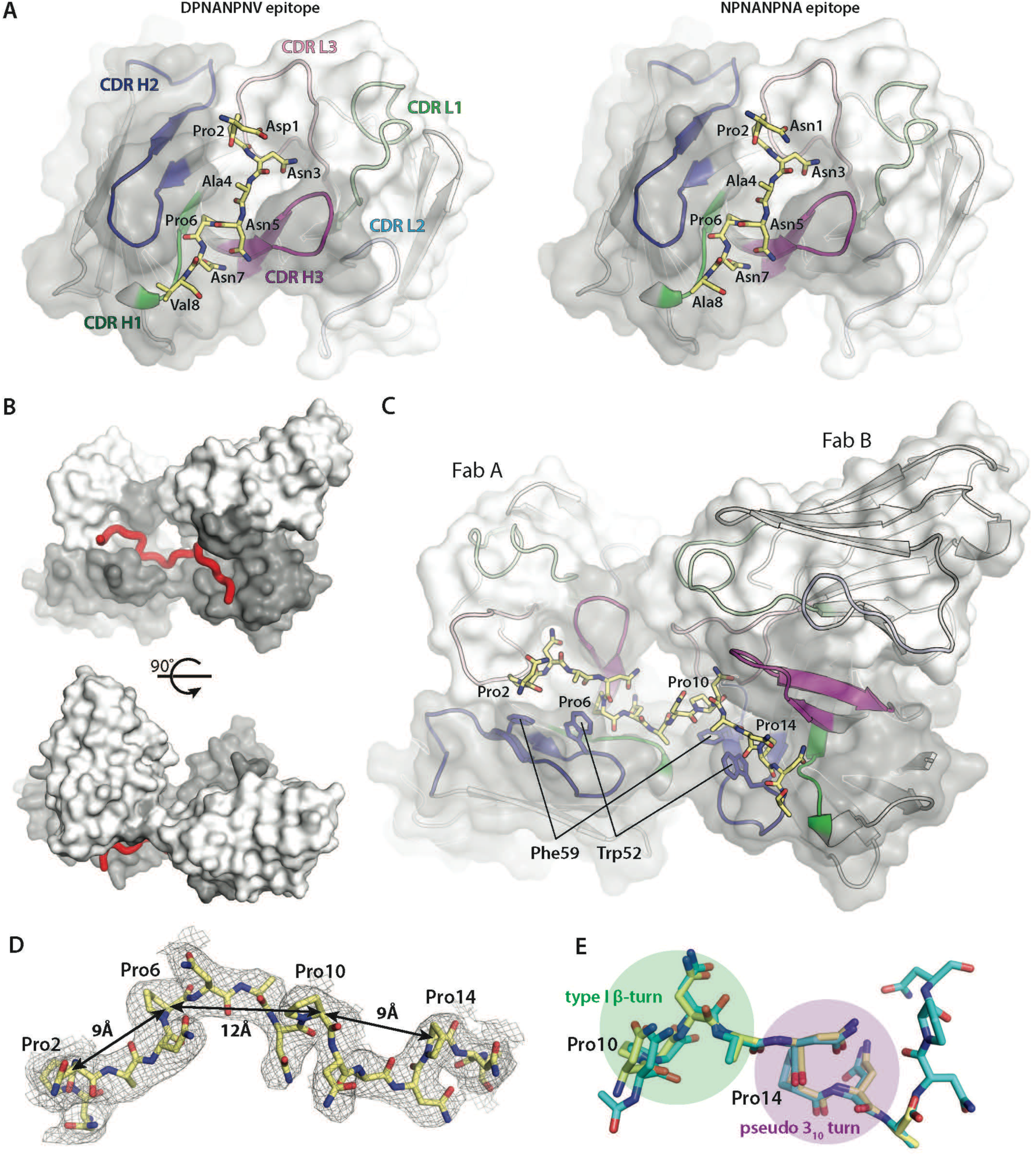
Epitope of two adjacent Fabs from the cryo-EM structure. (**A**) Illustration of the two different epitopes, DPNANPNV and (NPNA)_2_, that are observed in the cryo-EM rsCSP-Fab311 model, The heavy and light chain surfaces of the Fab variable domains are shown in gray and white, respectively. The three-dimensional structures of the variable domains are shown in cartoon representation with CDR H1, H2, H3, L1, L2, L3 colored in green, blue, magenta, light green, cyan and pink respectively. The rsCSP repeats are shown in stick representation (yellow carbons) and their amino acids are labeled and numbered from 1-8. (**B**) Front and side view of the composite epitope on the variable domains of two Fabs, labeled A and B, that bind two adjacent (NPNA)_2_ repeats, with the tetrapeptide shown as a red tube. (**C**) Detailed overview of the epitope shown in **B**. The tetrapeptide (NPNA) _4_ is shown as sticks (yellow carbons) with the 4 prolines labeled as Pro2, Pro6, Pro10 and Pro14, with the (NPNA) _4_ repeats numbered 1 to 16. The hydrophobic residues, Trp52 and Phe29, that engage in CH/*π* interactions with the repeat‘s proline residues, are shown as sticks. (**D**) The cryo-EM density for the (NPNA) _4_ repeats is contoured at 5 σ and shown as a gray mesh. Distances between the C α atoms of consecutive proline residues are highlighted. (**E**) Overlay of the peptide epitope from the cryo-EM structure (yellow carbons) with the corresponding peptide from the Fab311-(NPNA)_3_ crystal structure (teal carbons, PDB 6AXK) show minimal differences. The type I *β*-turn and pseudo 3_10_ turn are highlighted by transparent green and magenta circles respectively.

Helical conformations for NANP repeats have been proposed for CSP using computational methods. Gibson *et al.* used a modified buildup procedure to explore possible helical conformations for the (NANP) _6_ peptide, assuming that tandem repeats are likely to display helical or near-helical conformations driven by cooperative interactions (21). Two lowest energy helices were identified, with a radius of 3.5Å and 3.7Å, and a pitch of 10Å and 7Å respectively. Brooks *et al.* proposed an alternate much wider 12_38_ helix with a pitch of 4.95Å using molecular dynamics (MD) calculations (22). Interestingly, another model previously suggested a stem-like superhelix for the complete NANP repeat region with a width of 15 Å (radius 7.5 Å), length of 180 Å, and pitch of 7 NPNA repeats (23). Each of these helical predictions are very different from our structure. We observe a wider radius of 13.4 Å, length of 145 Å, and a much larger pitch of 49 Å (9.5 NPNA repeats, Fig. 1,D and F). To complete a full turn on the spiral, a fifth Fab partially packs underneath the first Fab, thereby making the pitch similar to the width of Fab311 along the longitudinal axis of the spiral (44.7 Å (24)). Our structure also differs from a recent model for an anti-NANP mouse antibody, 2A10, as a complex with NANP repeats. Here, antibodies are proposed to bind a narrow helix of repeats that adopt type I β-turns derived from MD simulations (25).

### Fab311 epitope on rsCSP

Traditionally, the repeat region of PfCSP has been described by the number of NANP repeats. However, NMR and X-ray crystallographic evidence show that the repeats are likely organized as NPNA structural motifs (14-18, 19). Hence, we adopt the NPNA nomenclature when discussing the epitope instead of the more general NANP notation. The Fab311 epitope was proposed to consist of a minimum of two to three NPNA repeats based on the crystal structure of Fab311 with the (NPNA)_3_ peptide and Isothermal Titration Calorimetry (ITC) affinity measurements (14). Here, the cryo-EM structure determines unambiguously that the epitope consists of only two NPNA repeats. In fact, the Fabs are so closely packed against one another that their two epitopes are seamlessly stitched together without the need of an additional repeat as a spacer. Furthermore, we observed that Fab311 is able to bind the NVDP repeats, thereby increasing the available epitopes on rsCSP from 15 (NPNA only) to 22 (including the DPNA and NPNV repeats). Remarkably, the only two sequence differences in DPNANPNV from NPNANPNA occurs on the edge of the epitope and thus are likely minimally inhibitory to Fab311 binding (Fig. 2). The Asp at the N-terminus is in a similar conformation to the Asn and the Val projects out into solvent. The calculated buried surface area (BSA) when taking two adjacent Fabs as one binding unit is 972Å^2^on the Fabs and 843Å2 on the (NPNA)_4_ peptide. Fabs are positioned such that the groove in which the peptide resides extends from one Fab directly into the other (Fig. 2, A and B). Overall, there is excellent agreement with the epitope in the cryo-EM structure with the first two of the three NPNA repeats observed in the crystal structure (Fig. 2E). The two repeats of the NPNA epitope adopt a type I *β*-turn followed by a pseudo 3_10_ turn (Fig. 2D) that repeats throughout the length of the spiral structure. Each pseudo 3_10_ turn has its asparagine (i) sidechain hydrogen bonding with the backbone amide of the next asparagine (i+2). Due to this unique repetition of the (NPNA)_2_ epitope in rsCSP, the proline residues consistently point away from the center of the spiral, serving as anchor points to which the Fabs latch on (Fig. 1F). Notably, CH/π interactions of the prolines with Trp52 and Phe59 alternate (Fig. 2B), with Cα-Cα distances of 9Å and 12Å between each consecutive proline pair (Fig. 2C). Trp52 provides key contacts with the peptide and may account for the frequent selection of germline VH3-33 (and related VH3-30) for recognition of the NANP repeats (16-17, 26).

### Inter-Fab contacts stabilize the CSP spiral structure

It is unlikely that free PfCSP is predominantly present as a well-defined spiral on the surface of the PfSPZ, since the repeat region is predicted to be disordered (27), and Atomic Force Microscopy (AFM) and single-molecule Force Microscopy experiments indicate that PfCSP can adopt multiple conformations (28, 29). Thus, binding of Fab311 may induce and stabilize the rigid spiral structure in the NANP repeat region of PfCSP. Surprisingly, neighboring Fabs that bind adjacent (NPNA)_2_ epitopes contribute 319Å^2^and 340Å^2^BSA to a novel interface between the Fabs (Fig. 3B). Taking into account these additional contacts, the total BSA on each Fab with rsCSP and neighboring Fabs becomes 1145Å^2^((972Å^2^/2)+319Å^2^+340Å^2^), which increases the original Fab-peptide BSA more than 2-fold. Close inspection reveals that the inter-Fab BSA between two Fabs (A and B) binding successive epitopes of the rsCSP (interface 1) spiral consists of polar contacts that are made between ^B^CDR L3/^A^CDR H3 and ^B^CDR H2/^A^CDR H1 (Fig. 3C). Interestingly, many residues that are involved in inter-Fab contacts correlate with affinity maturation from the IGHV3-33*01 and IGLV1-40*01 germline genes for the heavy and light chain, respectively (Fig. 3, G and H). Notably, salt bridges are made between Asp99 of ^A^CDR H3 and Arg93 and Arg94 of ^B^CDR L3; additionally, a cation-*π* interaction is found between Arg94 of ^B^CDR L3 and Tyr98 of ^A^CDR H3 where Arg94 N*ε* and the center of the aromatic tyrosine ring are 4.2Å apart (Fig. 3D). Furthermore, Asn31 of ^A^CDR H1 and Arg56, Asn57 and Glu64 of ^B^CDR H2 form an extensive hydrogen bonding network, which would be abrogated if reverted to the germline sequence (Fig. 3E). Most of these residues do not contact the NPNA repeat motifs, except for Asn31. Affinity maturation of Ser31 to Asn31 is likely driven by inter-Fab contacts, since Asn31 hydrogen bonds with the repeats using its main-chain atoms, while simultaneously forming a hydrogen bond with a neighboring Fab using its side chain (Fig. 3F). Other Fabs in close proximity are those that bind four epitopes away (B and F) such that they complete a full spiral turn and are either above or underneath the Fab of interest. Although some BSA is present between these two Fabs (interface 2), there are no direct contacts as assessed by CONTACSYM (Fig. 3A).

**Fig. 3.**
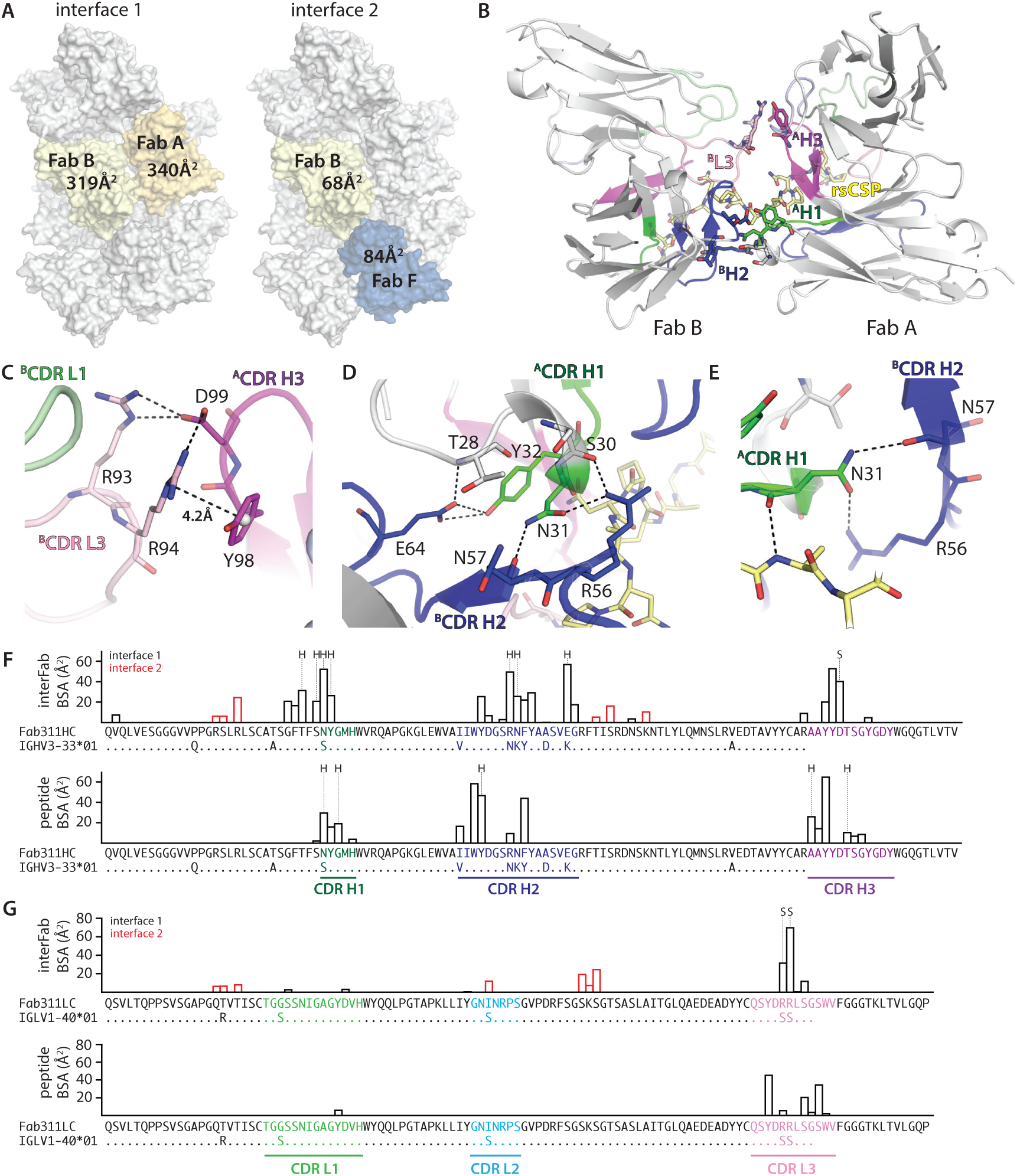
Affinity-matured inter-Fab contacts. (A) Surface representation of the rsCSP-Fab311 cryo-EM structure. Only the variable domains of Fab311 are shown. The Fabs of interest for which their mutual buried surface area (BSA) was calculated are colored yellow or blue, while other Fabs are colored gray. (B) Interactions between two Fabs (A and B) that bind adjacent epitopes on rsCSP. The Fabs are shown in cartoon representation with their heavy and light chains colored dark gray and light gray, respectively, and their CDR H1, H2, H3, L1, L2, L3 colored green, blue, magenta, light green, cyan and pink respectively. The tetrarepeat (NPNA)_4_ is colored yellow and shown in a cartoon representation. The CDR loops that engage in inter-Fab contacts are highlighted. (**C**) Salt bridges in interface 2 between Arg93 and Arg94 of ^B^CDR L3 (pink) and Asp99 of ^A^CDR H3 (magenta), and a cation-*π* interaction between Arg94 of ^B^CDR L3 and Tyr98 of ^A^CDR H3 are shown as black dashed lines. Tyr98 is V_H_ also shown because of its large BSA contribution. (**D**) Hydrogen bonding network (black dashed lines) in interface 2 between residues of ^B^CDR H2 and ^A^CDR H1. (**E**) Asn31 of ^A^CDR H1 hydrogen bonds with Ala8 of (NPNA)_4_ using its main-chain carbonyl (identical numbering as Figure 2C), while its side chain hydrogen bonds with the backbone of Asn57 and the side chain of Arg56 of ^B^CDR H2 from a neighboring Fab. (**F, G**) Individual residue contributions to the BSA of inter-Fab and to the peptide repeat contacts are shown in a bar plot for the heavy chain (**F**) and light chain (**G**). The CDRs as defined by Kabat are colored as in the previous figures. In addition, affinity-matured residues in V_H_ and V_L_ are shown by alignment of the Fab311 sequence with the germline V_H_ and V_L_ gene sequences (not including CDR H3). Residues that engage in hydrogen bonding and salt bridges are marked with ‘H’ or ‘S’ respectively.

### Mutagenesis of the Fab311 interface

To investigate the specificity of the interactions between adjacent Fabs, affinity-matured residues that engage with neighboring Fabs were mutated to the inferred germline sequence (Fab311 inter-Fab contact residue reverted, Fab311R). Specifically, four and two residues were mutated in the heavy (N31S, R56N, N57K and E64K) and light (R93S and R94S) chains, respectively. First, we assessed whether Fab311R can still bind to the (NPNA)_2_ peptide using ITC affinity measurements (fig. S3, table S2) and found that its binding is unperturbed indicating that few if any mutations are required for high affinity peptide binding. Next, we determined if the germline reversion mutagenesis abrogated formation of the rsCSP spiral using nsEM. Surprisingly, the 2D class averages revealed a new phenotype with varying stoichiometries for the rsCSP-Fab311R complex in which a well-defined long-range spiral was absent. Nevertheless, the rsCSP-Fab311R particles still adopted curved conformations in which the Fabs can still bind relatively closely together, indicating that some form of inter-Fab contacts may be encoded in the germline (Fig. 4). Such heterogeneity led to the inability of the particles to converge into a stable 3D reconstruction, which could not be further refined. By comparison, 2D class averages of wild-type rsCSP-Fab311 complex show a much more homogeneous and compact complex, providing further evidence that the affinity-matured inter-Fab residues play a crucial role in stabilizing the spiral architecture of rsCSP and presumably help gain increased avidity to CSP.

**Fig. 4.**
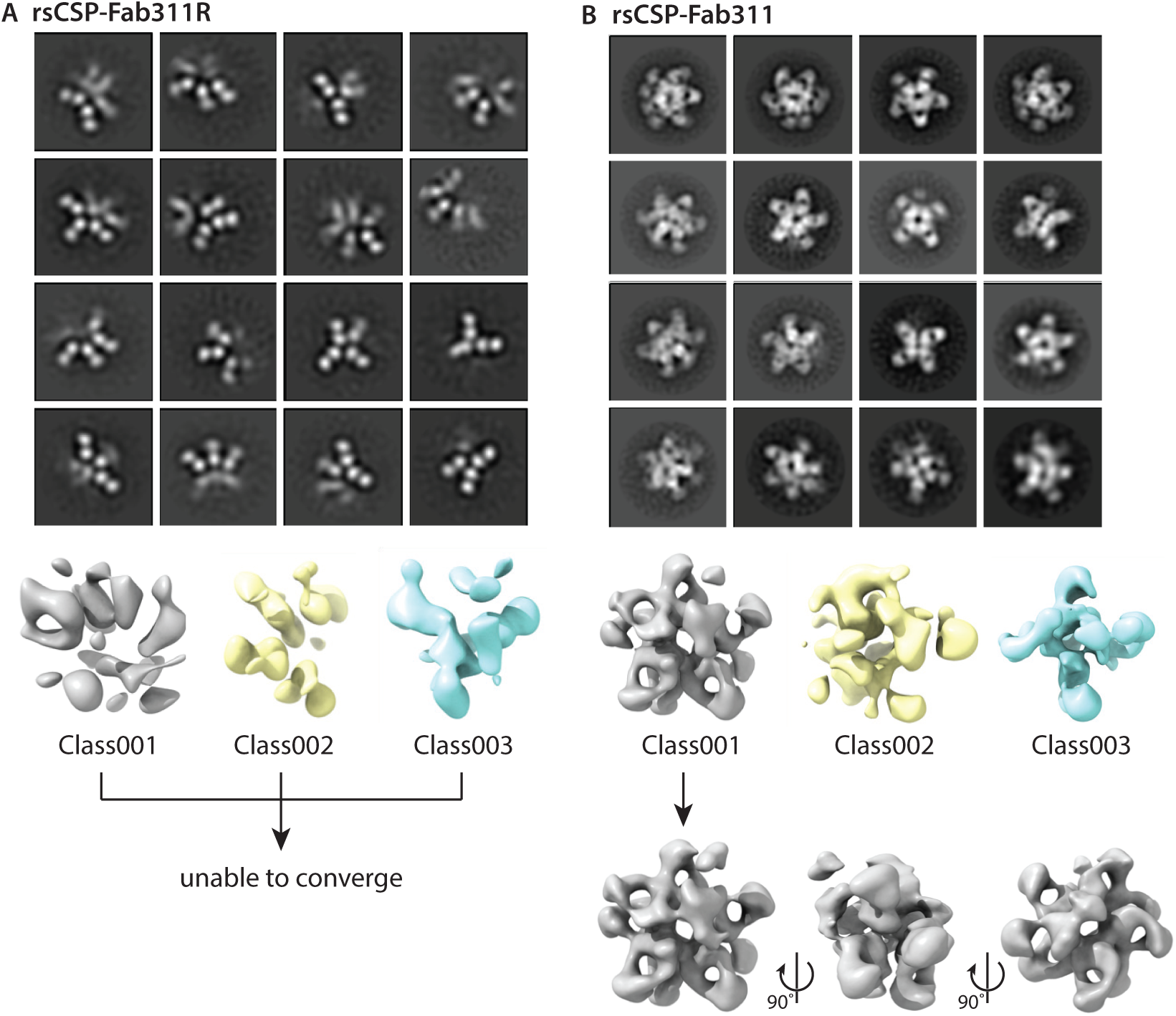
Germline reversion of inter-Fab contact residues. Representative class averages for (**A**) the germline-reverted Fab311 (Fab311R) in complex with rsCSP versus (**B**) full affinity-matured Fab311 in complex with rsCSP (14). 3D classes of rsCSP-Fab311R (1-3) did not converge during refinement, while class001 of the rsCSP-Fab311 complex converged and could be further refined.

### Conservation of the spiral architecture

To visualize how Fab311 might bind to the more physiologically relevant PfCSP on the PfSPZ surface, we expressed a full-length PfCSP (flCSP, based on the 3D7 strain) for nsEM studies with Fab311. The amino-acid sequence of flCSP is identical to rsCSP with the exception of the repeat region, which has 38 NANP repeats and 4 NVDP repeats, of which three are located at the N-terminus and one in the middle of the NANP repeat region (Fig. 1E). The 3D reconstruction of flCSP-Fab311 revealed an identical helical architecture to the rsCSP-Fab311 complex (Fig. 5A, figs. S4 and S5). Since the number of NANP repeats is doubled in flCSP compared to rsCSP, we were expecting >20 bound Fabs. However, the total Fab count in the flCSP-Fab311 complex is only 14. One possible explanation is that the additional NVDP repeat in the center of the NANP repeat region breaks up the NPNA registry and rigidity of the structure, since the affinity for NVDP is approximately 5-fold less than for NANP repeats (14). Nonetheless, these results provide evidence that the spiral architecture can also be formed by PfCSP with a widely different repeat length.

**Fig. 5.**
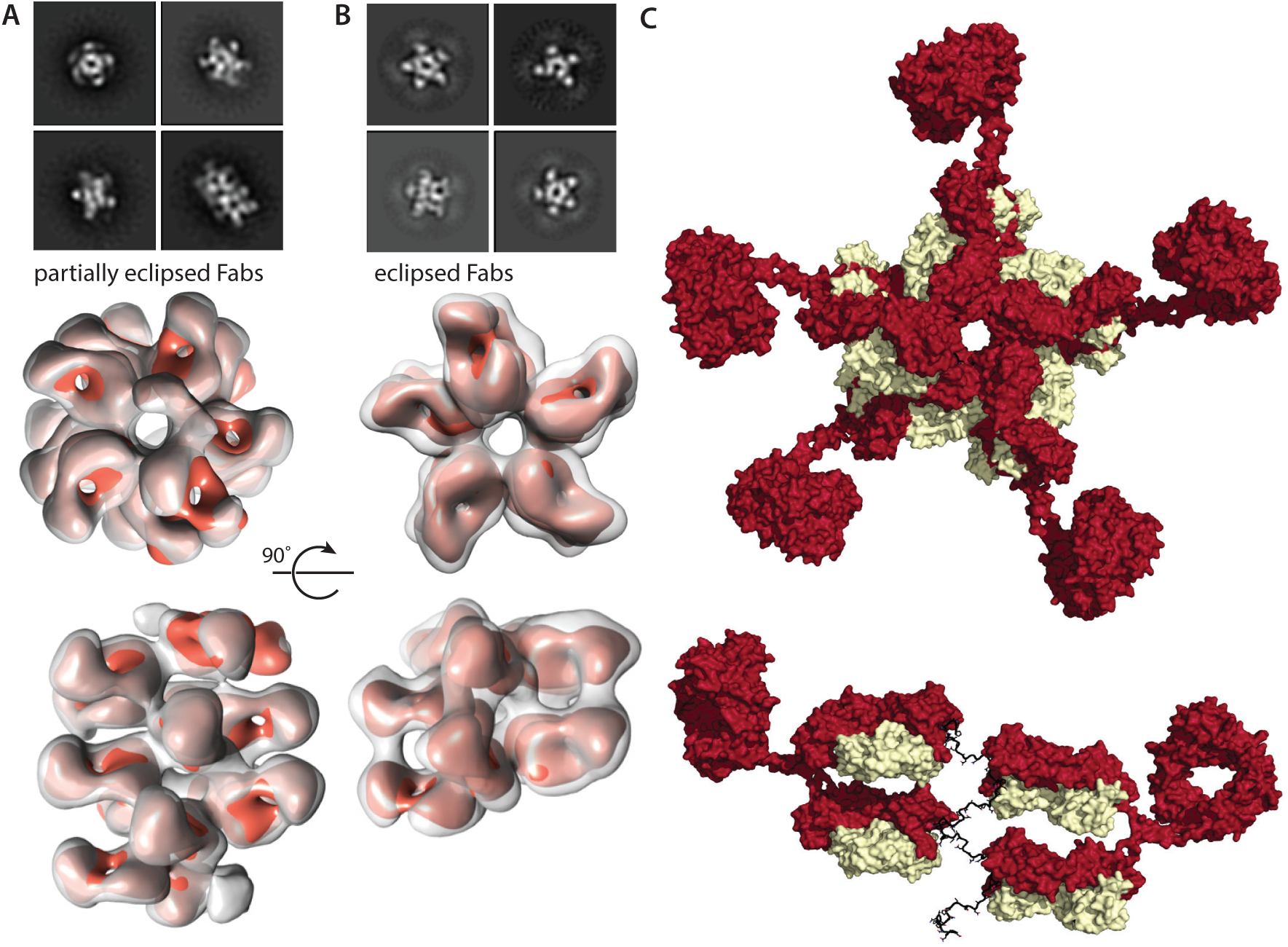
Helical architecture of full-length CSP with Fab311 and rsCSP in the presence of IgG311. Select reference-free 2D class averages and corresponding number of 30 Å low pass filtered Fab311‘s (orange) were docked into the nsEM 3D reconstructions of (**A**) flCSP in complex with Fab311 or (**B**) rsCSP in complex with IgG311. In both instances, CSP remains helical, with one complete turn equaling 5 epitopes. 10 Fabs (5 IgG311‘s) can bind to rsCSP, with the Fabs assigned to one IgG molecule eclipsing each other (when viewed down the helical axis) compared to the partially eclipsed orientation of the Fabs in the flCSP-Fab311 complex. (**C**) A model of the rsCSP-IgG311 complex shows that the Fabs of IgG311 bind rsCSP parallel with respect to each other, above and below each other on the spiral, but the Fabs from one IgG are separated on a linear scale along the spiral by 4 intervening Fabs. The Fc region was added to these two Fabs and the structure energy minimized with Rosetta.

To answer the question of whether an individual IgG is capable of binding to two epitopes within the same rsCSP molecule and further stabilize the spiral, we prepared a rsCSP-IgG311 complex for nsEM studies. A significant amount of aggregation was observed upon addition of IgG version of mAb311 (IgG311) to rsCSP as a result of crosslinking rsCSP molecules, which has also been termed the CSP reaction (30). After removal of aggregates by spin filtration (0.22 m) and subsequent size-exclusion chromatography (SEC), we were able to separate the sample into soluble aggregates, rsCSP-IgG311 complex, and unbound IgG311 fractions (Fig. S4). In the nsEM 2D classes of the rsCSP-IgG311 complex, the Fc domains appeared as diffuse densities radiating from the Fabs that did not converge in the 3D reconstruction (Fig. 5B). The 3D reconstruction closely matched the nsEM map of rsCSP-Fab311, but with a subtle difference in the helical twist. Comparison of the top views of the two reconstructions shows that Fab311 binds rsCSP in partially eclipsed orientations along the length of the spiral, while the two Fab domains of each bound IgG311 lie on top of one another (Fig. 5A and B, figs. S4 and S5). Notwithstanding, the rsCSP still adopts a spiral structure of identical radius with the IgG, despite the additional geometric constraints that the hinge region of the IgG311 poses on binding. A total of 5 IgG‘s (10 Fabs), were bound to rsCSP, in comparison to only 9 Fab311 in the nsEM 3D reconstruction (14). Thus, although IgG311 likely crosslinks PfCSP on the surface of PfSPZs, analysis of this minor population of single particles indicates that, just as for Fab311, IgG311 can bind with its Fab arms closely together and then still accommodate inter-Fab domain contacts between two different IgG molecules. The two Fabs that contribute to one IgG then are oriented such that the heavy and light chains are arranged light-heavy_ light-heavy and are not symmetric (light-heavy_heavy-light) as depicted in cartoons in most text books (Fig. 5C).

### Structural ramifications and implications for vaccine design

The cryo-EM reconstruction of rsCSP saturated with Fab311 at 3.4 Å demonstrates an unprecedented open spiral structure of rsCSP, which is still present with IgG or with flCSP. This structure differs substantially from previous predicted helical models for the NANP repeat region. Unexpectedly, the Fab domains not only make specific interactions with the NANP repeat region, but also with neighboring Fabs along the NANP spiral surface. These inter-Fab contact residues have undergone somatic hypermutation and are crucial for spiral formation. This finding provides strong evidence for antigen-induced maturation of inter-Fab interactions for human antibodies, which may prove to be a common mechanism for increasing affinity against the PfCSP repeat region and for tandem repeat sequences in general. Recently, heavy chain antibody fragments (nanobodies) derived from Alpacas against a pentameric antigen were observed to have inter-nanobody contacts, suggesting that this mechanism may be present across certain antibodies in different animal kingdoms (31). A previous structure of antibody 2G12 to HIV Env revealed a novel domain swap within the Fabs of a single IgG molecule, where the heavy chain from one Fab paired with the light chain of the other Fab, such that a new V_H_-V_H_ interface was formed that was also subject to affinity maturation (32). However, that configuration differs from the Fab arrangement here, where we observe instead affinity maturation between the Fabs that are connected to different IgG molecules. We do not know whether spiral formation correlates with protection, since mAb317 is bound in a less regular way to rsCSP, while being of similar efficiency as mAb311 in reducing the parasite liver load in mice experiments (14) (33) (fig. S6). Interestingly however, mAb317 2D class averages of the Fab bound to rsCSP are topologically similar to the more ordered mAb311 classes. Thus, it is likely that parts of the spiral may be present in the PfCSP conformational ensemble, perhaps even in the form of successive type-I β and pseudo 3_10_ turns, which in effect may code for the spiral preference in the presence of Fab311. If protection is correlated with recognition of a particular conformation of the PfCSP repeat region (20), inter-Fab maturation and spiral formation could lead to higher avidity and potentially more protective anti-malaria antibodies.

A recently described human antibody (MGG4) bound to the N-terminal junction peptide (KQPADGNPDPNANP) showed binding to an NPDP repeat in the junction region just prior to the repeat region (16), which Fab311 is also capable based on our cryoEM structure. Additionally, Fab311 and MGG4 have identical heavy-chain germline gene (VH3-33/30) usage and mode of binding through CH/*π* interactions between a proline in a pseudo 3_10_ turn with conserved Trp52 (Fig. 1a). Since Fab311 is derived from a volunteer immunized with RTS,S and MGG4 from a volunteer immunized with irradiated PfSPZs, the similarities between the two imply that the previously reported potent public antibody lineage, from which MGG4 originates (16), can be accessed using the RTS,S vaccine candidate. This intriguing cryoEM structure may provide the basis for design of previously unanticipated novel immunogens that now can take into account the three-dimensional spiral architecture of the CSP repeat region rather than information derived solely from Fab-peptide studies with smaller numbers of repeats.

## ACKNOWLEDGEMENTS

We thank B. Anderson for maintaining the microscopes and H.L. Turner, C.A. Bowman, and G. Ozorowski for technical assistance. We thank Kelsey Mertes and Ashley Birkett of PATH MVI for critical reading and comments on the manuscript. This work was funded by PATH‘s Malaria Vaccine Initiative and the Bill and Melinda Gates Foundation (grant no. OPP1170236) under collaborative agreements with The Scripps Research Institute.

## REFERENCES AND NOTES

1. World Health Organization, World malaria report 2016 (World Health Organization, Geneva, 2016).

2. M. Imwong, T. T. Hien, N. T. Thuy-Nhien, A. M. Dondorp, N. J. White, Spread of a single multidrug resistant malaria parasite lineage (PfPailin) to Vietnam. Lancet Infectious Diseases 17, 1022–1023 (2017).

3. R. S. Nussenzweig, V. Nussenzweig, Development of sporozoite vaccines. Philos Trans R Soc Lond B Biol Sci 307, 117–128 (1984).

4. F. Zavala et al., Rationale for development of a synthetic vaccine against *Plasmodium falciparum* malaria. Science 228, 1436–1440 (1985).

5. N. M. Bowman et al., Comparative population structure of *Plasmodium falciparum* circumsporozoite protein NANP repeat lengths in Lilongwe, Malawi. Sci Rep 3, 1990 (2013).

6. D. E. Neafsey et al., Genetic diversity and protective efficacy of the RTS, S/AS01 malaria vaccine. N Engl J Med 373, 2025–2037 (2015).

7. J. B. Ancsin, R. Kisilevsky, A binding site for highly sulfated heparan sulfate is identified in the N terminus of the circumsporozoite protein: significance for malarial sporozoite attachment to hepatocytes. J Biol Chem 279, 21824–21832 (2004).

8. M. B. Doud et al., Unexpected fold in the circumsporozoite protein target of malaria vaccines. Proc Natl Acad Sci U S A 109, 7817–7822 (2012).

9. M. De Wilde, J. Cohen. (Google Patents, 2001).

10. RTS, S Clinical Trials Partnership et al., First results of phase 3 trial of RTS, S/AS01 malaria vaccine in African children. N Engl J Med 365, 1863–1875 (2011).

11. RTS, S Clinical Trials Partnership et al., Efficacy and safety of the RTS, S/AS01 malaria vaccine during 18 months after vaccination: a phase 3 randomized, controlled trial in children and young infants at 11 African sites. PLoS Med 11, e1001685 (2014).

12. RTS, S Clinical Trials Partnership et al., Efficacy and safety of RTS, S/AS01 malaria vaccine with or without a booster dose in infants and children in Africa: final results of a phase 3, individually randomised, controlled trial. Lancet 386, 31–45 (2015).

13. K. A. Collins, R. Snaith, M. G. Cottingham, S. C. Gilbert, A. V. S. Hill, Enhancing protective immunity to malaria with a highly immunogenic virus-like particle vaccine. Sci Rep 7, 46621 (2017).

14. D. Oyen et al., Structural basis for antibody recognition of the NANP repeats in *Plasmodium falciparum* circumsporozoite protein. Proc Natl Acad Sci U S A 114, E10438–E10445 (2017).

15. G. Triller et al., Natural parasite exposure induces protective human anti-malarial antibodies. Immunity 47, 1197–1209.e10 (2017).

16. J. Tan et al., A public antibody lineage that potently inhibits malaria infection through dual binding to the circumsporozoite protein. Nat Med 24, 401–407 (2018).

17. N. K. Kisalu et al., A human monoclonal antibody prevents malaria infection by targeting a new site of vulnerability on the parasite. Nat Med 24, 408–416 (2018).

18. H. J. Dyson, A. C. Satterthwait, R. A. Lerner, P. E. Wright, Conformational preferences of synthetic peptides derived from the immunodominant site of the circumsporozoite protein of Plasmodium falciparum by 1H NMR. Biochemistry 29, 7828–7837 (1990).

19. A. Ghasparian, K. Moehle, A. Linden, J. A. Robinson, Crystal structure of an NPNA-repeat motif from the circumsporozoite protein of the malaria parasite *Plasmodium falciparum*. Chem Commun, 174–176 (2006).

20. J. A. Regules et al., Fractional third and fourth dose of RTS, S/AS01 malaria candidate vaccine: A phase 2a controlled human malaria parasite infection and immunogenicity study. J Infect Dis 214, 762–771 (2016).

21. K. D. Gibson, H. A. Scheraga, Predicted conformations for the immunodominant region of the circumsporozoite protein of the human malaria parasite Plasmodium falciparum. Proc Natl Acad Sci U S A 83, 5649–5653 (1986).

22. B. R. Brooks, R. W. Pastor, F. W. Carson, Theoretically determined three-dimensional structure for the repeating tetrapeptide unit of the circumsporozoite coat protein of the malaria parasite *Plasmodium falciparum*. Proc Natl Acad Sci U S A 84, 4470–4474 (1987).

23. M. L. Plassmeyer et al., Structure of the *Plasmodium falciparum* circumsporozoite protein, a leading malaria vaccine candidate. J Biol Chem 284, 26951–26963 (2009).

24. ^L^Gly68 and ^H^Ser74 C_α_-C_α_ distance

25. C. R. Fisher et al., T-dependent B cell responses to *Plasmodium* induce antibodies that form a high-avidity multivalent complex with the circumsporozoite protein. PLoS Pathog 13, e1006469 (2017).

26. R. Murugan et al., Clonal selection drives protective memory B cell responses in controlled human malaria infection. Sci Immunol 3, eaap8029 (2018).

27. A. J. Guy et al., Insights into the immunological properties of intrinsically disordered malaria proteins using proteome scale predictions. PLoS One 10, e0141729 (2015).

28. R. Herrera et al., Reversible conformational change in the *Plasmodium falciparum* circumsporozoite protein masks its adhesion domains. Infect Immun 83, 3771–3780 (2015).

29. A. P. Patra, S. Sharma, S. R. Ainavarapu, Force spectroscopy of the Plasmodium falciparum vaccine candidate circumsporozoite protein suggests a mechanically pliable repeat region. J Biol Chem 292, 2110–2119 (2017).

30. J. Vanderberg, R. Nussenzweig, H. Most, Protective immunity produced by the injection of x-irradiated sporozoites of Plasmodium berghei. V. In vitro effects of immune serum on sporozoites. Mil Med 134, 1183–1190 (1969).

31. R. A. Bernedo-Navarro et al., Structural basis for the specific neutralization of Stx2a with a camelid single domain antibody fragment. Toxins (Basel) 10, E108 (2018).

32. D. A. Calarese et al., Antibody domain exchange is an immunological solution to carbohydrate cluster recognition. Science 300, 2065–2071 (2003).

33. Because of the unique spiral architecture of rsCSP when bound to Fab311, we collected a cryo-EM dataset on another protective antibody, Fab317, in complex with rsCSP to improve the resolution of the previously published nsEM map thereby gaining molecular details. Briefly, Fab317 was also isolated from the phase IIa RTS, S/AS01B CHMI clinical trial, has the identical germline gene as mAb311 (VH3-33/30), and provided 99.7% reduction of parasite liver load, as previously described (14). Fab317 binds up to 3 NANP repeats in comparison to Fab311 which only requires 2 repeats. However, inspection of the 2D class averages revealed various stoichiometries similar to the nsEM 2D classes, with up to five Fab317’s bound to rsCSP. Although the dataset was subject to extensive rounds of computational processing and 3D sorting, the particles were unable to converge to high resolution (fig. S6).

